# An inhibitor of apoptosis (SfIAP) interacts with SQUAMOSA promoter binding protein (SBP) transcription factors that exhibit pro-cell death characteristics

**DOI:** 10.1101/267435

**Authors:** Ryan Kessens, Nicholas Sorensen, Mehdi Kabbage

## Abstract

Despite the functional conservation of programmed cell death (PCD) across broad evolutionary distances, an understanding of the molecular machinery underpinning this fundamental program in plants remains largely elusive. This is despite its critical importance to development, homeostasis, and proper responses to stress. Progress in plant PCD has been hindered by the fact that many core regulators of animal PCD are absent in plant genomes. Remarkably, numerous studies have shown that the ectopic expression of animal anti-PCD genes in plants can suppress cell death imposed by many stresses. In this study, we capitalize on the ectopic expression of an insect inhibitor of apoptosis (SfIAP) to identify novel cell death regulators in plants. A yeast two-hybrid assay was conducted using SfIAP as bait to screen a tomato cDNA library. This screen identified several transcription factors of the SQUAMOSA promoter binding protein (SBP) family as potential SfIAP binding partners. We confirmed this interaction *in vivo* for our top two interactors, SlySBP8b and SlySBP12a, using coimmunoprecipitation. Interestingly, overexpression of *SlySBP8b* and *SlySBP12a* induced spontaneous cell death in *Nicotiana benthamiana* leaves. Overexpression of these two transcription factors also induced the accumulation of reactive oxygen species and enhanced the growth of the necrotrophic pathogen *Alternaria alternata*. Fluorescence microscopy confirmed the nuclear localization of both SlySBP8b and SlySBP12a, while SlySBP12a was also localized to the ER membrane. These results support a pro-death role for SlySBP8b and SlySBP12a and provide potential targets that can be utilized to improve stress tolerance in crop plants.

**Highlights:** SBP transcription factors SlySBP8b and SlySBP12a from tomato interact with an insect inhibitor of apoptosis protein (SfIAP). Both exhibit pro-cell death characteristics while SlySBP12a activity may be regulated through ER membrane tethering.

**Abbreviations:**

PCD: programmed cell death
IAP: inhibitor of apoptosis
BIR: baculovirus IAP repeat
RING: really interesting new gene
FB1: fumonisin B1
SBP: SQUAMOSA promoter binding protein
ROS: reactive oxygen species
35S: cauliflower mosaic virus 35S promoter
HA: hemagglutinin
YFP: yellow fluorescent protein
DAB: 3,3’-Diaminobenzidine
QIS-Seq: quantitative interactor screen sequencing
CLSM: confocal laser scanning microscopy
DHE: dihydroethidium
NLS: nuclear localization signal
TMD: transmembrane domain
ER: endoplasmic reticulum
HR: hypersensitive response
MTTF: membrane-tethered transcription factor

## Introduction

Programmed cell death (PCD) is a fundamental aspect of development and stress response that is conserved throughout all kingdoms of life (Allocati *et al.*, 2015). This process of genetically controlled cellular suicide has been studied extensively in animal systems, and the results of these research efforts have led to major treatment advances for many human diseases (Fuchs and Steller, 2011). In contrast, our understanding of the biochemical pathways underlying PCD in plants is severely lacking. This is largely due to the absence of obvious orthologs of core regulators of apoptosis, a well-studied form of PCD in animals (Kabbage *et al.*, 2017). While this has undoubtedly slowed progress on plant PCD research, it has also presented a unique opportunity for the discovery of novel regulators of PCD in plant systems.

Apoptosis is a specific type of PCD characterized by distinct morphological and biochemical features (Kroemer *et al.*, 2009). Apoptotic cell death in animals is executed through the activation of <u>c</u>ysteine-dependent <u>asp</u>artate-specific prote<u>ases</u> termed caspases. Caspases exist as inactive pro-enzymes that can be activated by external or internal cellular cues. Once activated, caspases execute an orderly demise of the cell by targeting negative regulators of apoptosis, cytoskeletal components, and other caspases (Parrish *et al.*, 2013). Due to the terminal nature of apoptosis, caspases must be kept under multiple layers of regulation. The Inhibitor of Apoptosis (IAP) family is an important group of proteins that negatively regulate caspase activity. The defining feature of all IAPs is the presence of one or more Baculovirus IAP Repeat (BIR) domains, which confer substrate specificity (Verhagen *et al.*, 2001). Additionally, some IAPs contain a Really Interesting New Gene (RING) domain that serves as a functional E3 ubiquitin ligase domain. Inhibitor of Apoptosis proteins can inhibit caspase activity by preventing pro-caspases from becoming active or by suppressing active caspases. This can be accomplished by simply blocking the active site pocket of a caspase or by utilizing the RING domain to ubiquitinate a caspase and mark it for proteasome-mediated degradation (Feltham *et al.*, 2012; Gyrd-Hansen and Meier, 2010).

Despite the fact that obvious orthologs of IAPs and caspases are absent in plant genomes, the ectopic expression of animal and viral apoptotic regulators in tobacco (*Nicotiana* spp.) and tomato (*Solanum lycopersicum*) modulate plant cell death. This was first reported nearly two decades ago when the expression of *Bax*, a mammalian pro-apoptotic gene absent in plant genomes, induced localized tissue collapse and cell death in *Nicotiana benthamiana* (Lacomme and Santa Cruz, 1999). Shortly thereafter, Dickman *et al.* (2001) demonstrated that expression of a viral *IAP*(*OpIAP*), as well as anti-apoptotic members of the Bcl-2 family, conferred resistance to a suite of necrotrophic fungal pathogens in *Nicotiana tabacum*. Pathogens with a necrotrophic lifestyle require dead host tissue for nutrient acquisition and studies on *Cochliobolus victoriae, Sclerotinia sclerotiorum,* and *Fusarium* spp. revealed that these necrotrophic fungal pathogens hijack host cell death machinery to kill cells (Asai *et al.*, 2000; Glenn *et al.*, 2008; Kabbage *et al.*, 2013; Lorang *et al.*, 2012; Williams *et al.*, 2011).

More recently, we showed that overexpression of an *IAP* from *Spodoptera frugiperda* (fall armyworm; *SfIAP*) in tobacco and tomato prevented cell death associated with a wide range of abiotic and biotic stresses (Kabbage *et al.*, 2010; Li *et al.*, 2010). Tobacco and tomato lines expressing *SfIAP* had increased heat and salt stress tolerance, two abiotic stresses that induce cell death. These transgenic lines were also resistant to the fungal necrotroph *Alternaria alternata* and the mycotoxin fumonisin B1 (FB1) (Li *et al.*, 2010). Fumonisin B1 is produced by some species of *Fusarium* and is a potent inducer of apoptosis in animal cells and apoptotic-like PCD in plant cells (Gilchrist, 1997).

It has been over 15 years since it was first reported that overexpression of animal anti-apoptotic regulators in plants conferred enhanced resistance against a wide assortment of necrotrophic pathogens. During this time, numerous studies have confirmed the efficacy of animal apoptotic regulators in plants without identifying the means by which these regulators function. In this study, we used an unbiased approach to identify *in planta* binding partners of SfIAP in tomato to better understand how this insect IAP is able to inhibit cell death and confer stress tolerance in plants. Yeast two-hybrid and coimmunoprecipitation (CoIP) assays show that SfIAP interacts with members of the SQUAMOSA promoter binding protein (abbreviated SBP in tomato or SPL in some other species) transcription factor family. Overexpression of two tomato SBPs, *SlySBP8b* and *SlySBP12a*, induced cell death in tobacco leaves accompanied by enhanced production of reactive oxygen species (ROS). Overexpression of *SlySBP8b* and *SlySBP12a* also created an environment that was more conducive to the growth of the necrotrophic fungal pathogen *A. alternata*. In summary, our findings uncover SlySBP8b and SlySBP12a as novel SfIAP binding partners that exhibit pro-death attributes.

## Materials and Methods

### Plant material and growth conditions

*Nicotiana benthamiana* plants were grown on a 16 h light cycle (~50 microeinsteins m^−2^ s^−1^) at 26° C and ~60% humidity. *Nicotiana glutinosa* (PI 555510) and tomato (*Solanum lycopersicum* cv. Bonny Best) plants were grown on a 16 h light cycle (~100 microeinsteins m^−2^s^−2^) at 22° C and ~60% humidity. The soil composition for all plants consisted of SunGro^®^ propagation mix and Sunshine^®^ coarse vermiculite in a 3:1 ratio. Plants were watered with deionized water supplemented with Miracle-Gro^®^ all-purpose fertilizer (1g/L) as needed.

### Plasmid construction

The full-length open reading frames of *SlySBP-like* (Solyc07g062980), *SlySBP4* (Solyc07g053810), *SlySBP6a* (Solyc03g114850), *SlySBP6c* (Solyc12g038520), *SlySBP8b* (Solyc01g090730), and *SlySBP12a* (Solyc01g068100) were amplified by PCR from cDNA collected from tomato inflorescence tissue (Supplemental Table S1). AttB1 and attB2 adapters were added to forward and reverse primers, respectively, to generate attB-flanked amplicons suitable for Gateway^TM^ Recombination Cloning (Invitrogen).

Amplicons were recombined into the entry vector pDONR™/Zeo using BP clonase II (Invitrogen). *SlySBP8b(NLS_mt_)* and *SlySBP12a(NLS_mt_)* constructs were generated using the Q5^®^ Site-Directed Mutagenesis Kit (New England Biolabs). *SlySBP12a(ΔTMD)* and *TMD*_*SlySBP12a*_ were amplified from *SlySBP12a* in pDONR™/Zeo using the primers indicated in Supplementary Table S1 and recombined into pDONR™/Zeo. For overexpression in *N. benthamiana* leaves and tomato protoplasts, entry vectors were mixed with the desired pEarleyGate destination vectors (Earley *et al.*, 2006) and recombined using LR clonase II (Invitrogen). pEarleyGate vectors drive transgene expression using a cauliflower mosaic virus 35S (35S) promoter and were used to generate N-terminal yellow fluorescent protein (YFP; pEarleyGate104) or N-terminal influenza hemagglutinin (HA; pEarleyGate201) fusions. All constructs were verified using Sangar sequencing before being transformed into *Agrobacterium tumefaciens* GV3101.

Plasmids for the yeast two-hybrid screen were prepared as follows. *SfIAP*, *SfIAP*_*BIR1*_, and *luciferase* cDNAs were cloned into the bait vector pGilda under control of the *GAL1* promoter and in-frame with an N-terminal fusion of the *E. coli* LexA DNA binding protein (Takara Bio USA, Inc.). *Luciferase* (firefly luciferase from *Photinus pyralis*) was cut from an existing plasmid using a 5’-Nco1 restriction site in the START codon and a 3’-Not1 restriction site outside of the ORF and ligated into pGilda. Primers for *SfIAP* (GenBank: AF186378.1) and *SfIAP*_*BIR1*_ amplification were designed to place an EcoR1 site at the 5’ end and a BamH1 site at the 3’ end of the ORF. Primers used for amplification can be found in Supplementary Table S1. Amplicons were cut using these restriction enzymes and ligated into pGilda. Tomato cDNAs were expressed from the *GAL1* promoter with an N-terminal fusion of the B42 activation protein in the pB42AD plasmid (Takara Bio USA, Inc.). Bait and prey library were sequentially transformed into EGY48 yeast using standard protocols.

### Yeast two-hybrid screening

Yeast containing bait and plasmid were plated on SD galactose (-His/-Trp/-Leu) to induce gene expression and select for bait-prey interactions. After incubating at 28°C for ~5 days, colonies were pooled in 10 mL of sorbitol/phosphate buffer (1.2 M sorbitol, 0.1 M NaPO_4_, pH 7.5) per plate, pelleted, and resuspended in 2 mL of sorbitol/phosphate buffer supplemented with 500 U of lyticase (Sigma: L2524-25KU) and 250 μg of RNAse A. Yeast cells were incubated in the lyticase buffer for 3 h at 37°C prior to plasmid recovery. Plasmid DNA was extracted using a Wizard Plus SV Miniprep kit (Promega) and a modified protocol. Briefly, 2.5 mL of lysis solution and 80 uL of alkaline protease solution were added to yeast protoplasts and incubated at room temperature for 10 mins. Next, 3.5 mL of neutralization solution was added and cellular debris was pelleted by centrifugation. Supernatant was run through the provided columns and plasmid DNA eluted according to the manufacturer’s instructions. Low-cycle PCR was performed to amplify cDNA’s from the prey library. Briefly, MyFi™ proofreading DNA polymerase (Bioline) and pB42AD forward and reverse primers (flanking the cDNA insertion site of pB42AD) were used to amplify cDNA’s (Supplementary Table S1). A QIAquick PCR purification kit (Qiagen) was used to clean PCR products before sequencing.

### Illumina sequencing and data analysis

Sequencing was performed by the Biotechnology Center at UW-Madison using Illumina Next Generation sequencing with 100 bp paired-end reads. The sequencing data were uploaded to the Galaxy web platform, and we used the public server at *usegalaxy.org* to analyze the data (Afgan *et al.*, 2016). Reads were groomed and trimmed to remove low quality bases and adapter sequences before alignment (Bolger *et al.*, 2014). Bait (pGilda) and prey (pB42AD) plasmid sequences were concatenated with the *Saccharomyces cerevisiae* reference genome (S288C_reference_sequence_R64-2-1_20150113) to create a FASTA file containing sources of plasmid and gDNA contamination. Reads were aligned to this file using Bowtie 2 (Langmead and Salzberg, 2012). Aligned reads (plasmid and gDNA) were discarded while unaligned reads were aligned to the tomato reference genome (Solgenomics: ITAG2.4) with Bowtie 2. Cufflinks (v2.2.1) was used to assemble transcripts from these aligned reads and calculate FPKM values for each locus (Trapnell *et al.*, 2012). Enrichment scores for each locus were calculated using R Studio and scripts written in-house (RStudio Team, 2016). Details of Galaxy pipeline, in-house scripts, and complete dataset are available upon request.

### Transient expression in N. benthamiana and N. glutinosa

Agrobacterium strain GV3101 was grown overnight in liquid LB supplemented with gentamycin and kanamycin (50 μg/mL) at 28 ◻C with shaking. Cells were harvested by centrifugation, washed once with sterile deionized water, and resuspended in infiltration medium (10 mM MgSO_4_, 9 mM MES, 10 mM MgCl_2_, 300 μM acetosyringone, pH 5.7) to a final concentration of OD_600_ = 0.9. Cultures were incubated at room temperature for 4 h before infiltration. *Nicotiana benthamiana* plants were infiltrated with a 1-mL needleless syringe at 4-5 weeks of age with the two youngest and easily infiltratable leaves being used. *Nicotiana glutinosa* plants were infiltrated at 5-6 weeks of age with a single leaf being used on each plant, typically corresponding to the 4^th^ or 5^th^ true leaf. Plants were transformed at different ages due to differences in rate of growth between the two species.

For total protein extraction, leaf tissue was frozen in liquid nitrogen and ground in 3x Laemmli buffer (10% β-mercaptoethanol). Samples were boiled for 10 minutes followed by centrifugation at 10,000 *g* for 5 min. Supernatants were removed and transferred to new tubes. Total proteins were separated by electrophoresis on a 12% Tris-Glycine-SDS polyacrylamide gel (BioRad). Proteins were transferred to a nitrocellulose membrane. Total protein was detected using Ponceau S stain. Epitope-tagged proteins were detected by probing with α-GFP (Cell Signaling 2955S) or α-HA (Cell Signaling 3724S) primary antibodies. The α-GFP antibody was detected using goat α-mouse IgG conjugated to horseradish peroxidase (HRP) (Cell Signaling 7076P2) while the α-HA antibody was detected using goat α-rabbit IgG conjugated to HRP (Cell Signaling 7074P2). Amersham™ ECL™ reagent (GE Life Sciences) was used to detect HRP-conjugated antibodies.

### Transient transfection of tomato protoplasts

Mesophyll protoplasts form tomato cotyledons were isolated from 10 day-old plants using the Tape Sandwich method (Wu *et al.*, 2009). A total of 6 μg of plasmid was used for each transfection with an equal ratio used for cotransfections. Transfections were performed using polyethylene glycol (PEG) as described previously (Yoo *et al.*, 2007). Protoplasts were used for imaging the day after transfection.

### Coimmunoprecipitation assays

Agrobacterium strains harboring free *35S:YFP* or *35S:YFP-SfIAP^M4^(I332A)* were coinfiltrated with strains harboring *35S:HA-SlySBP8b* or *35S:HA-SlySBP12a*. A 7:2 ratio of YFP strains to HA strains was used due to relatively low accumulation of YFP-SfIAP^M4^(I332A) protein compared to HA-SlySBP8b and HA-SlySBP12a. Approximately 40 h post-agroinfiltration, transformed leaves were collected and ground in liquid nitrogen to a fine powder. Extraction buffer (150 mM Tris-HCl, 150 mM NaCl, 5 mM EDTA, 0.2% IGEPAL, and 1% plant protease inhibitor cocktail [Sigma]) was added at a concentration of 2 mL/g of leaf tissue. YFP-tagged proteins were immunoprecipitated by incubating the lysate with α-GFP magnetic agarose beads (GFP-Trap_MA; Chromotek) for 2 h at 4° C. Beads were washed three times in extraction buffer (w/o IGEPAL) and boiled in 30 μL of 2x SDS loading buffer before loading on duplicate 12% Tris-Glycine-SDS polyacrylamide gels (BioRad). Proteins were transferred to duplicate nitrocellulose membranes and probed with α-GFP (Cell Signaling 2955S) or α-HA (Cell Signaling 3724S) primary antibodies. The α-GFP antibody was detected using goat α-mouse IgG conjugated to HRP (Cell Signaling 7076P2) while the α-HA antibody was detected using goat α-rabbit IgG conjugated to HRP (Cell Signaling 7074P2). Amersham™ ECL™ reagent (GE Life Sciences) was used to detect HRP-conjugated antibodies.

### Confocal laser scanning microscopy

Confocal laser scanning microscopy was performed on a Zeiss ELYRA LSM780 inverted confocal microscope using a 40x, 1.1-numerical aperture, water objective. YFP fusions, chlorophyll autofluorescence, and DHE were excited with a 488 nm argon laser. YFP emission was detected between 502-542 nm, chlorophyll emission was detected between 657-724 nm, and DHE was detected between 606-659 nm. mCherry was excited with a 561 nm He-Ne laser and emission was detected between 606-651 nm.

### Electrolyte leakage analysis

Cell death progression in *N. benthamiana* leaves was assessed by measuring ion leakage. Approximately 24 h post-agroinfiltration, eight leaf discs were collected from two leaves on the same plant and pooled into a single well of a 12-well plate. Leaf discs were washed for 30 min in 4 mL of deionized water by rotating plates at 50 rpm at room temperature. Wash water was removed and replaced with 4 mL of fresh deionized water. Immediately after adding fresh water, the conductivity of the solution was recorded, representing the 24 h post-agroinfiltration measurement. The conductivity of the water was measured using an ECTestr 11^+^ MultiRange conductivity meter (Oakton) at the indicated time points.

### DAB staining of N. benthamiana leaves

Staining solution was prepared by dissolving 3,3’-Diaminobenzidine (DAB; Sigma) in HCl at pH 2. Once dissolved, this solution was added to Na_2_HPO_4_ buffer (10 mM) for a final DAB concentration of 1 mg/mL. Tween-20 (0.05% v/v) was added and the final pH was adjusted to 7.2. Whole leaves were collected, placed in petri dishes, submerged in DAB staining solution, and vacuum infiltrated. Plates were covered in aluminum foil and incubated at room temperature with shaking. After 4 hours, DAB staining solution was removed and replaced with clearing solution A (25% acetic acid, 75% ethanol). Leaves were heated at 80◻C for 10 minutes to remove chlorophyll. Clearing solution A was removed and replaced with clearing solution B (15% acetic acid, 15% glycerol, 70% ethanol). Leaves were incubated in clearing solution B overnight at room temperature to remove residual chlorophyll.

### A. alternata inoculation of N. glutinosa leaves

*Alternaria alternata* isolated from potato was provided by Dr. Amanda Gevens (University of Wisconsin, Madison, WI). Leaves were harvested from *N. glutinosa* plants one day post-agroinfiltration. For FB1 treatments, leaves were coinfiltrated with an Agrobacterium suspension containing the *35S:YFP* construct and 5 μM FB 1 (Cayman Chemicals). Detached leaves were placed adaxial-side up in petri dishes (100 mm × 20 mm) containing 3 layers of wet filter paper. Five-mm-diameter agar plugs were collected from the edge of an actively growing fungal colony on potato dextrose agar. Leaves were wounded with a 1 mL pipette tip along the midrib and agar plugs were placed fungal-side-down on top of the wound. Inoculated leaves were kept at room temperature (~23◻C) for the duration of the experiment.

### Image acquisition and analysis

All leaf images were taken using a Nikon D5500 camera with a Nikon AF-S NIKKOR 18-55 mm lens. Quantification of DAB staining intensity and fungal growth was performed using the Fiji package for ImageJ (Schindelin *et al.*, 2012). For quantification of DAB staining intensity, the Colour Deconvolution package was used to isolate the DAB color channel for each DAB-stained leaf (Ruifrok and Johnston, 2001). Staining intensity caused by *35S:YFP* expression on the left half of each leaf was subtracted from the staining intensity caused by *35S:HA-SlySBP8b* or *35S:HA-SlySBP12a* expression on the right half of the same leaf. Fungal lesions were quantified by tracing the periphery of the lesion and calculating the area within the periphery using ImageJ. Statistical analyses were performed using a one-way analysis of variance (ANOVA) with Tuckey’s honest significant difference (HSD) test in R Studio (RStudio Team, 2016).

## Results

### Identification of SfIAP binding partners in tomato

To identify putative binding partners of SfIAP from tomato, we performed a yeast two-hybrid assay coupled with next-generation sequencing using a method developed by Lewis *et al.* (2012) termed quantitative interactor screen sequencing (QIS-Seq). This method enables the entire pool of interactors to be sequenced by pooling all yeast colonies together instead of individually sequencing each colony using Sangar sequencing. The high throughput nature of QIS-Seq proved useful for screening multiple baits, including a negative control, against the library as well as sequencing the entire cDNA library itself (Supplementary Fig. S1A.).

SfIAP contains two BIR domains and a C-terminal RING domain. The BIR1 domain and the RING domain are essential for complete SfIAP function in plants while the BIR2 domain is dispensable (Kabbage *et al.*, 2010). Full-length SfIAP and the BIR1 domain alone (SfIAP_BIR1_) were used as bait to screen a tomato cDNA library produced under stressed conditions. The SfIAP_BIR1_ construct was used to prolong transient interactions that can occur upon ubiquitination of substrates by the RING domain of full-length SfIAP. Luciferase served as a negative control to account for non-specific protein interactions and potential autoactivation of the selectable marker. The cDNA library itself was also sequenced to account for biases in transcript abundance.

Enrichment scores were calculated for each locus using the equation in Supplementary Fig. S1B. A total of 13 putative interactors with enrichment scores of 50 or higher were identified in our screen (Table 1). Interestingly, this list contained six members of the SQUAMOSA promoter binding protein (SBP) family of transcription factors. Based on enrichment scores, the top interactor with full-length SfIAP was SlySBP8b (95.7) while the top interactor with SfIAP_BIR1_ was SlySBP12a (98.7). Also present at lower enrichments were SlySBP4, -6a, -6c, and an unannotated homolog referred to as SlySBP-like (Table 1).

**Table 1:**
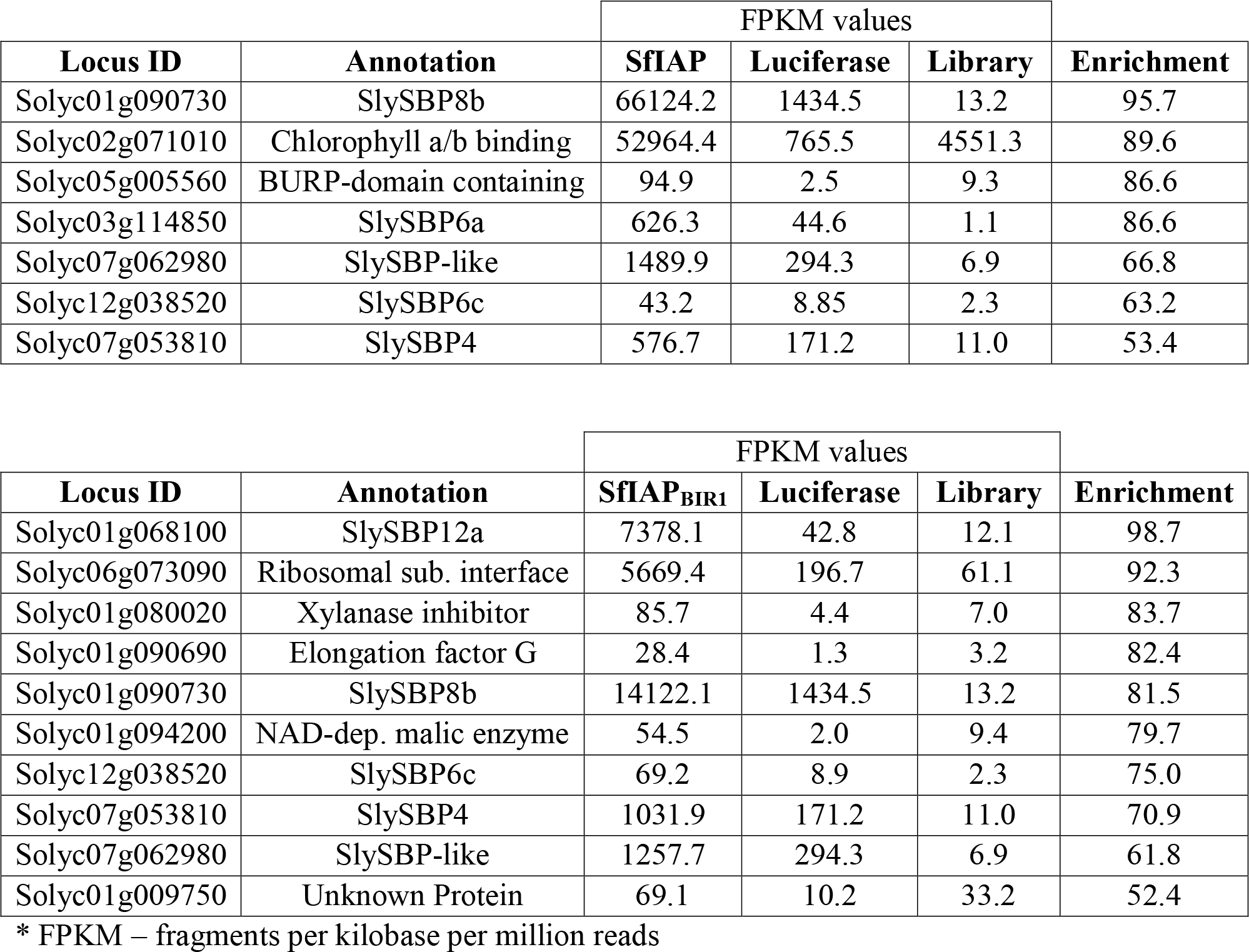
Enriched genes identified from QIS-Seq using full-length SfIAP or the BIR1 domain alone as bait.

### Induction of tissue death by SlySBP8b and SlySBP12a

SfIAP is known to inhibit apoptosis in *S. frugiperda* and suppress cell death when ectopically expressed in plants. Thus, we anticipated that SfIAP-interacting partners in plants may be positive regulators of cell death. To narrow our list of candidate genes, we transiently overexpressed cDNA clones of each tomato SBP identified from our yeast two-hybrid screen in *N. benthamiana* leaves and monitored these leaves for signs of tissue death. The generated cassettes contained an N-terminal hemagglutinin (HA) tag and were driven by a cauliflower mosaic virus 35S promoter (35S). At 5 days post-transformation, tissue collapse induced by *35S:HA-SlySBP8b* expression and a lesion-mimic phenotype induced by *35S:HA-SlySBP12a* expression could clearly be seen (Fig. 1A). However, overexpression of the other *SlySBP*s failed to produce any visible signs of tissue death. Immunoblots using an α-HA antibody confirmed protein accumulation for all constructs (Fig. 1B). These results show that at least two SfIAP interactors induce clear signs of cell death upon overexpression in *N. benthamiana*.

**Fig. 1.**
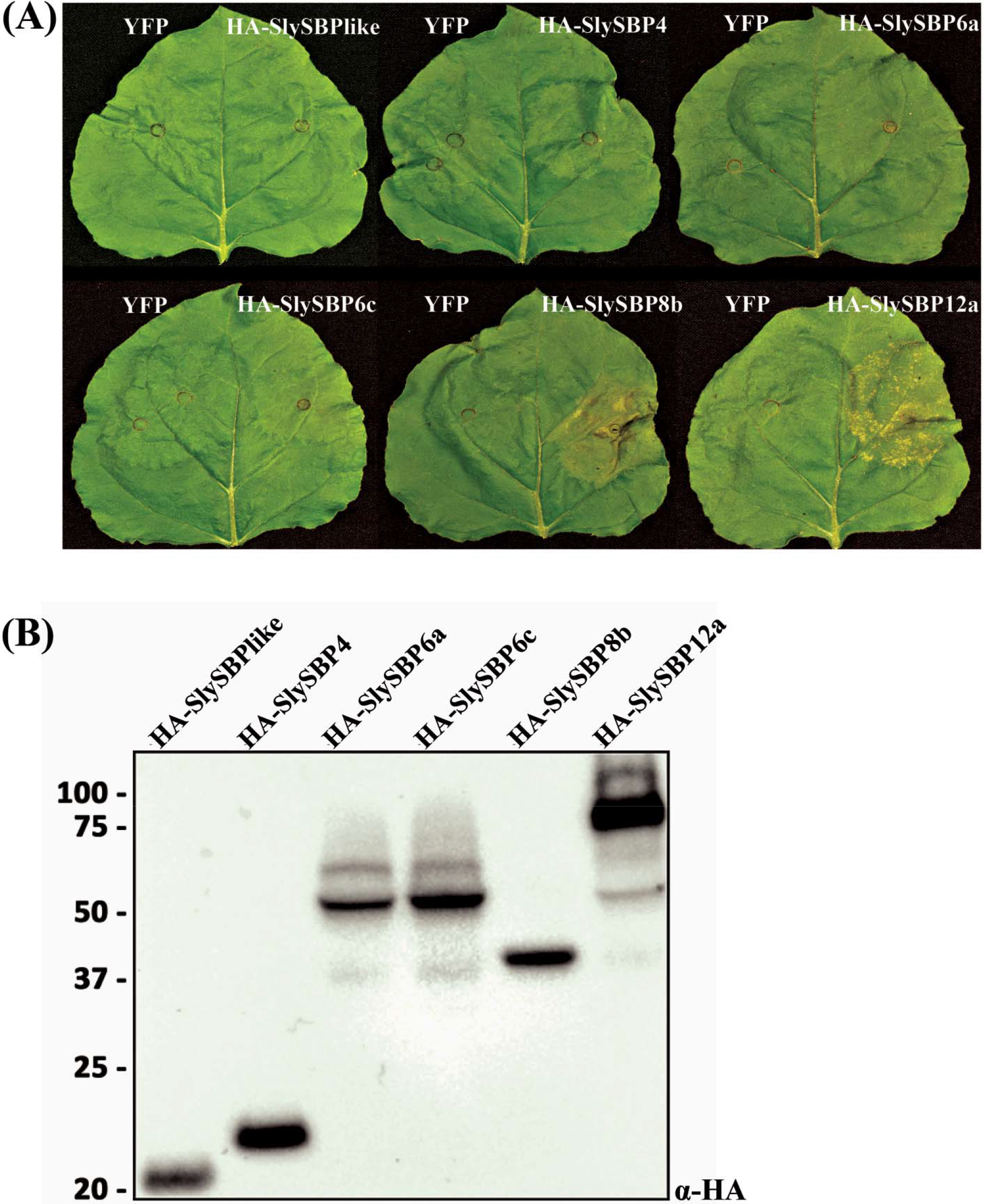
Cell death induced by overexpression of *SlySBP* transcription factors in *N. benthamiana*. Enriched SlySBP transcription factors from the yeast two-hybrid assay were transiently overexpressed in *N. benthamiana*. (A) The left half of each leaf was transformed with free *YFP* as a negative control while the right half was transformed with the corresponding *SlySBP* gene containing an N-terminal HA tag and 35S promoter. Images were taken 5 days post-transformation. (B) A Western blot was performed on tissue collected 2 days post-transformation to confirm accumulation of SlySBP proteins. Proteins were detected using an α-HA antibody.

### SlySBP8b and SlySBP12a interact with SfIAP^M4^(I332A) in-planta

Due to the strong tissue death phenotype associated with the overexpression of *SlySBP8b* and *SlySBP12a*, we focused our subsequent efforts on these two SBP variants. For *in vivo* confirmation of the yeast two-hybrid results, we performed coimmunoprecipitation (CoIP) assays in *N. benthamiana* leaves. A truncated version of SfIAP beginning at the 4^th^ methionine residue was used as our bait. This version maintains its function in *S. frugiperda* cells but lacks a caspase recognition site that is typically cleaved in *S. frugiperda* (Cerio *et al.*, 2010). This is particularly important since we show that cleavage at the N-terminus of the full-length protein occurs in *N. benthamiana*, thus removing N-terminal epitope tags (Supplementary Fig. S2). To prolong transient interactions that may take place between SfIAP and its targets following ubiquitination, an E3 ligase mutant of the truncated SfIAP protein was used by mutating a conserved residue in the RING domain (Cerio *et al.*, 2010). This construct, referred to as SfIAP^M4^(I332A), is resistant to N-terminal cleavage in *N. benthamiana* (Supplementary Fig. S2).

Two days after coexpression of *35S:YFP-SfIAP^M4^(I332A)* with *35S:HA-SlySBP8b* or *35S:HA-SlySBP12a*, total proteins were extracted from leaves and incubated with GFP-Trap_MA beads (Chromotek, Germany). All proteins were detected in the input fraction, and HA-SlySBP8b and HA-SlySBP12a were successfully pulled-down by YFP-SfIAP^M4^(I332A) but not by free YFP (Fig. 2). These data confirm the yeast two-hybrid results and demonstrate that SfIAP^M4^(I332A) interacts with SlySBP8b and SlySBP12a in plant cells.

**Fig. 2.**
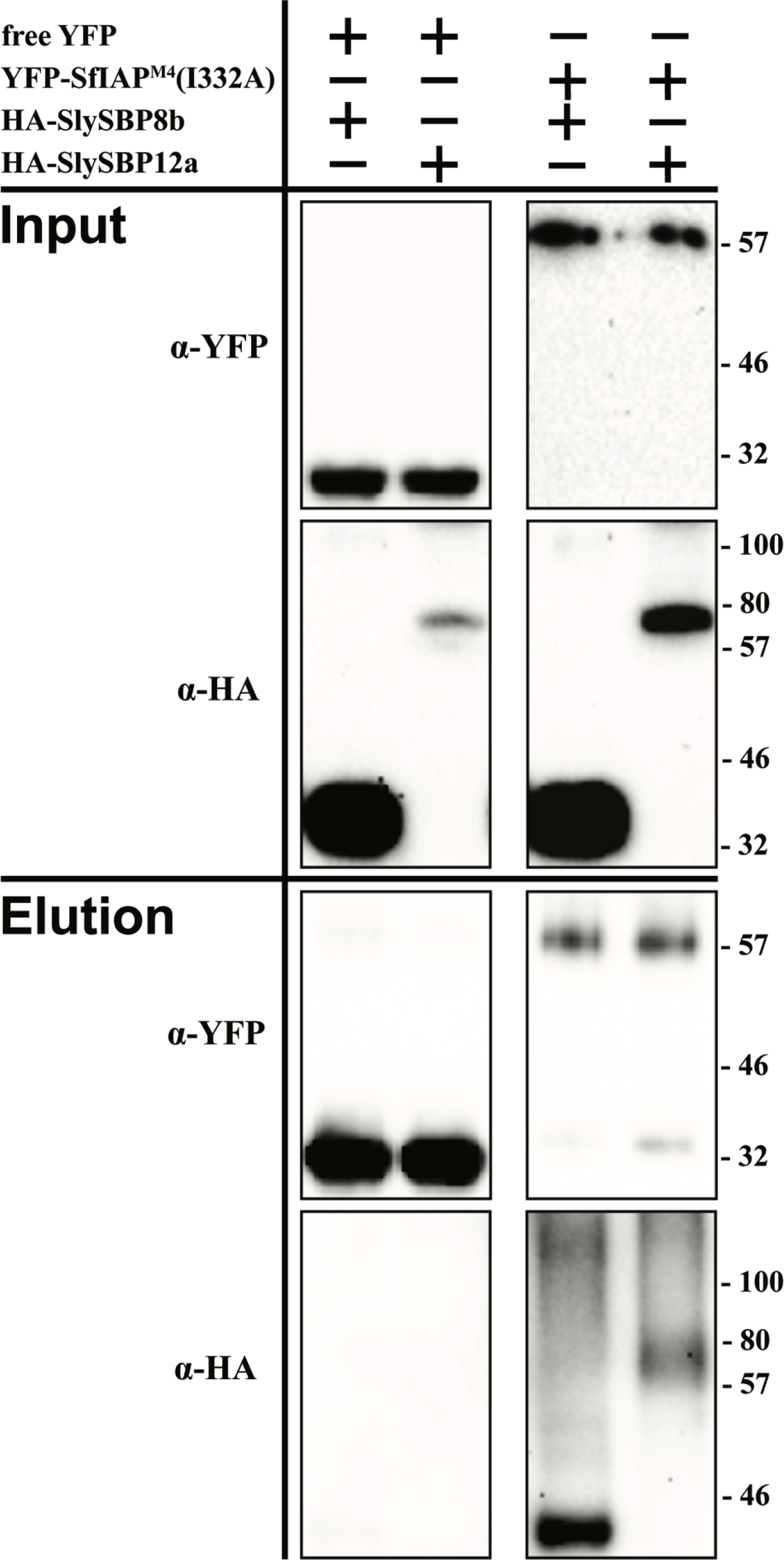
Coimmunoprecipitation of SflAP^M4^(I332A) with SlySBP8b and SlySBP12a in *N. benthamiana. 35S:YFP-SflAP^M4^(l332A)* or free *YFP* was transiently coexpressed with *35S:HA-SlySBP8b* or *35S:HA-SlySBP12a* in *N. benthamiana* leaves. Proteins were immunoprecipitated with an α-YFP affinity matrix. A portion of each sample was taken before immunoprecipitation to serve as the input control. An immunoblot was performed on input and elution fractions using the indicated antibodies to detect the epitope-tagged proteins.

### Role of SlySBP8b and SlySBP12a localization in cell death induction

As putative transcription factors, we reasoned that SlySBP8b and SlySBP12a function in the nucleus and nuclear localization would be required to regulate genes involved in cell death induction. Additionally, a predicted bi-partite nuclear localization signal (NLS) is present in the SBP domain of all tomato SBP transcription factors (Salinas *et al.*, 2012). Localization was assessed by expressing *35S:YFP-SlySBP8b* and *35S:YFP-SlySBP12a* in tomato mesophyll protoplasts. Confocal laser scanning microscopy (CLSM) revealed that both YFP-SlySBP8b and YFP-SlySBP12a colocalized with the nuclear marker dihydroethidium (DHE) (Fig. 3). To further substantiate the role of nuclear localization in cell death induction, site-directed mutagenesis was used to substitute conserved lysine and arginine residues in the NLS with leucine (Supplementary Fig. S3). Overexpression of the two NLS mutants, *35S:HA-SlySBP8b(NLS_mt_)* and *35S:HA-SlySBP12a(NLS_mt_)*, in *N. benthamiana* leaves did not induce visible signs of cell death (Fig. 4A). Immunoblots performed on tissues overexpressing both the wild-type and NLS mutants confirmed that protein accumulation was not greatly affected by mutations in the NLS (Fig. 4B). Thus, nuclear localization of these two transcription factors is required for cell death to occur.

**Fig. 3.**
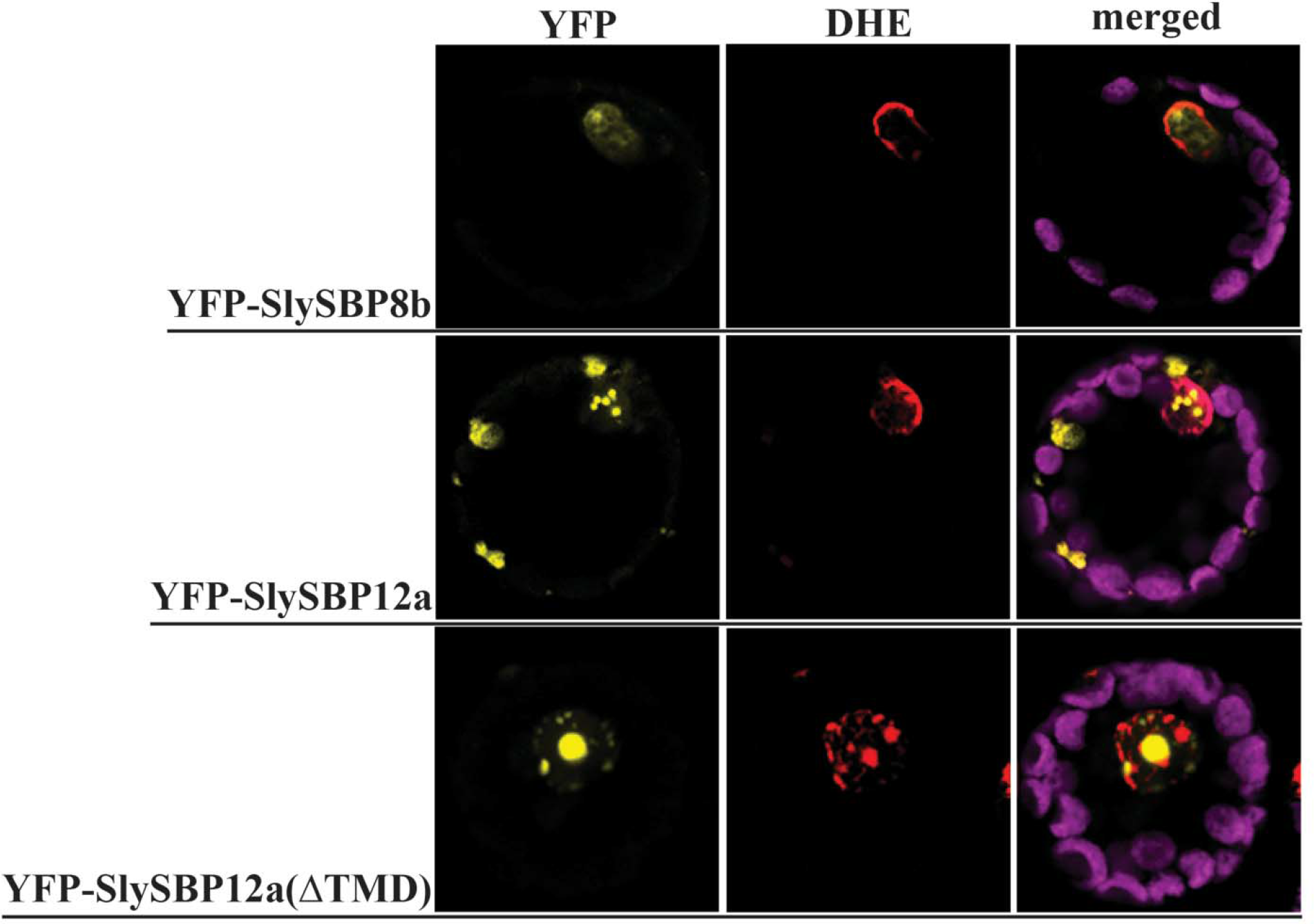
Nuclear localization of SlySBP8b, SlySBP12a, and SlySBP12a(ΔTMD) in tomato protoplasts. Tomato protoplasts were transfected with plasmids encoding *35S:YFP-SlySBP8b*, *35S:YFP-SlySBP12a*, or *35S:YFP-SlySBP12a(ΔTMD)* and imaged by CLSM. Dihydroethidium (red) was used as a nuclear counterstain while the magenta signal represents chloroplast autofluorescence.

**Fig. 4.**
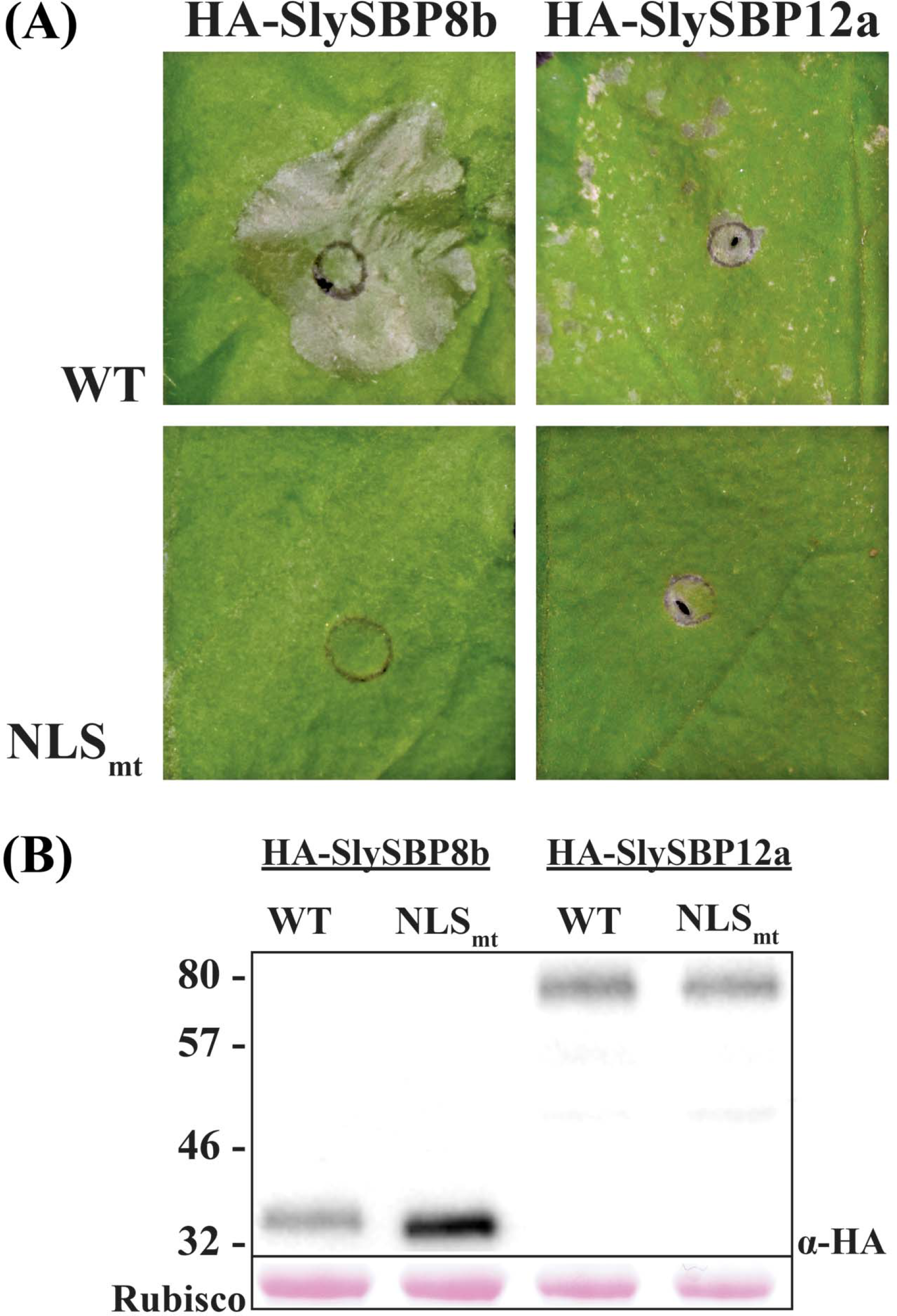
Disruption of the NLS in SlySBP8b and SlySBP12a prevents cell death in *N. benthamiana* upon overexpression. *35S:HA-SlySBP8b*, *35S:HA-SlySBP8b(NLS_mtl_)*, *35S:HA-SlySBP12a*, or *35S:HA-SlySBP12a(NLS_mt_)* were transiently transformed in *N. benthamiana*. (A) Images of leaves taken 5 days post-transformation. (B) Immunoblot performed on tissue collected 2 days post-transformation. An α-HA antibody was used to detect SlySBP proteins and Ponceau S stain was used to detect Rubisco as a loading control.

While YFP-SlySBP8b was found to be strictly nuclear-localized, YFP-SlySBP12a was also localized to diffuse pockets outside of the nucleus (Fig. 3). The presence of a putative C-terminal transmembrane domain (TMD) in SlySBP12a (Supplementary Fig. S3) suggested that this localization pattern could be due to the anchoring of SlySBP12a to a cellular membrane. Removal of the last 73 amino acids of SlySBP12a eliminated the putative TMD and resulted in complete localization of YFP-SlySBP12a(ΔTMD) to the nucleus (Fig. 3; Fig. 5). Additionally, overexpression of *35S:HA-SlySBP12a(ΔTMD)* in *N. benthamiana* caused enhanced cell death characterized by extensive tissue collapse at the site of transgene expression and increased electrolyte leakage compared to the full-length construct (Fig. 6A and 6B). The TMD of SlySBP12a may thus regulate its access to the nucleus and the subsequent induction of cell death.

**Fig. 5.**
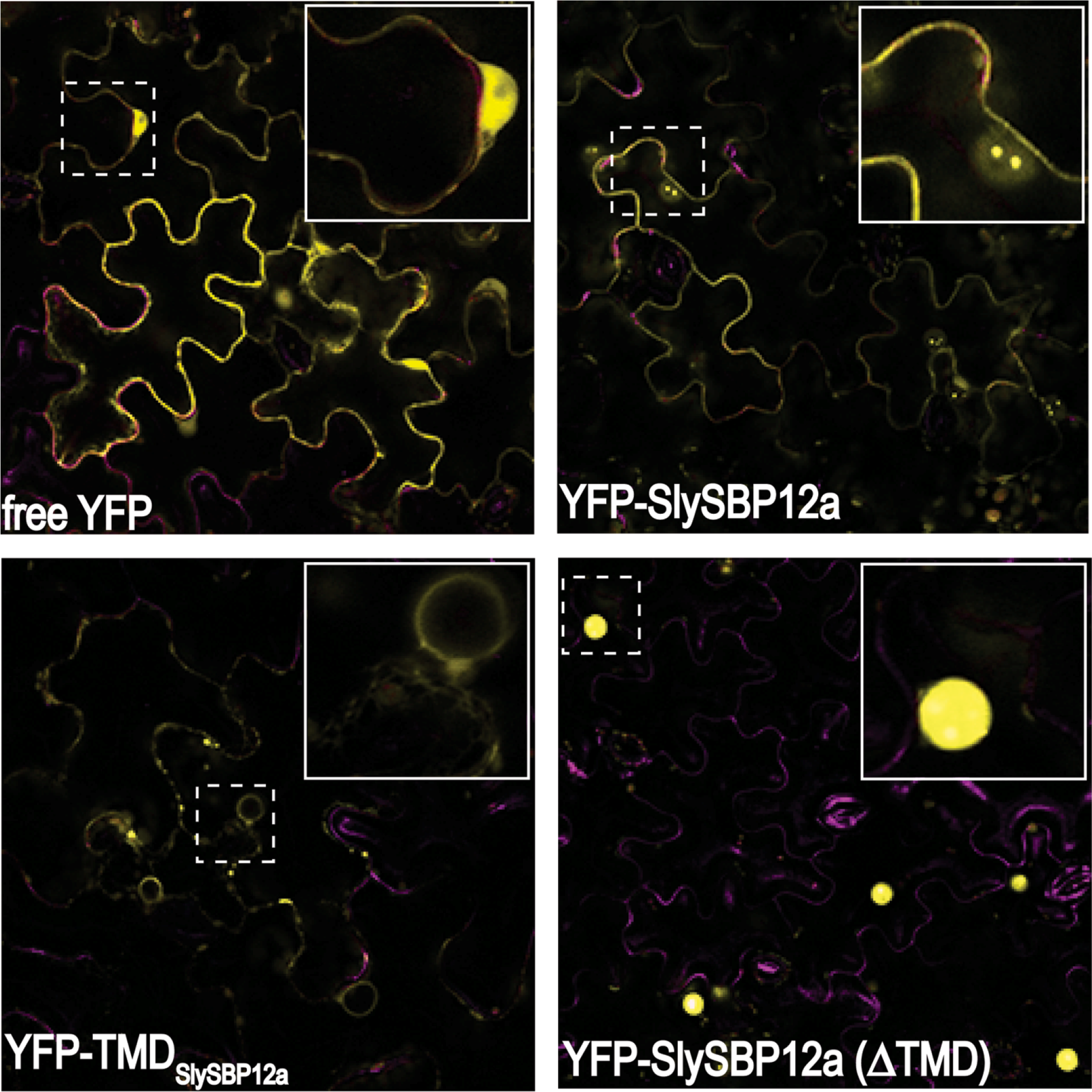
Localization of SlySBP12a, TMD_SlySBP12a_, and SlySBP12a(ΔTMD) in *N. benthamiana* epidermal cells. Leaves were transiently transformed with *35S:YFP-SlySBP12a*, *35S:YFP-SlySBP12a(ΔTMD)*, *35S:YFP-TMD*_*SlySBP12a*_, or *35S:YFP* and imaged using CLSM two days post-transformation. The dashed-line box in each panel is magnified and displayed in the upper-right corner of each panel. Chlorophyll autofluorescence is shown in magenta.

**Fig. 6.**
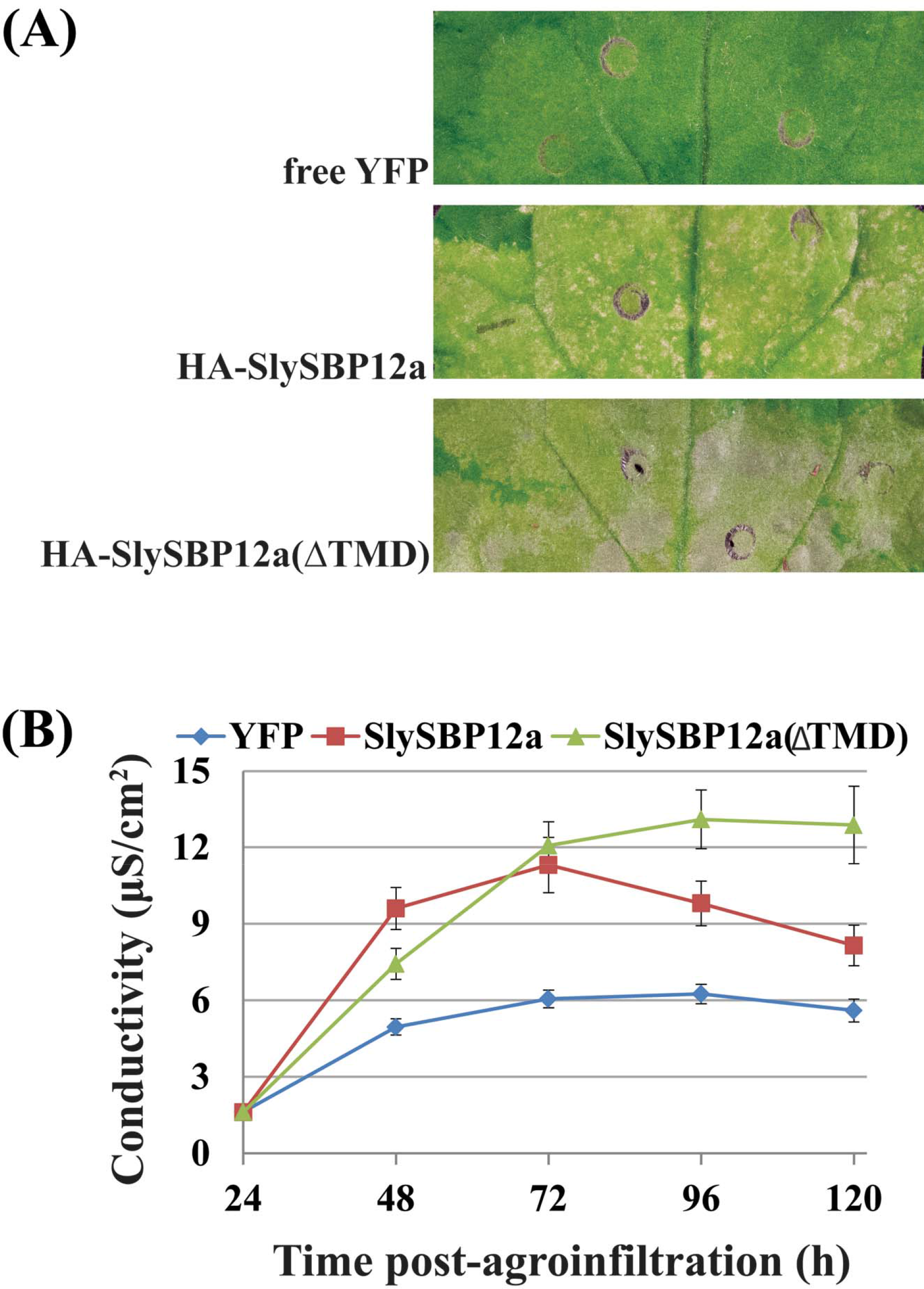
Removal of the TMD from SlySBP12a results in enhanced cell death upon overexpression. *35S:HA-SlySBP12a*, *35S:HA-SlySBP12a(ΔTMD)* or *35S:YFP* were transiently transformed in *N. benthamiana*. (A) Images of leaves taken 5 days post-transformation. (B) Electrolyte leakage assay used to quantify cell death. *35S:YFP* – blue diamond; *35S:HA-SlySBP12a* – red square; *35S:HA-SlySBP12a(ΔTMD)* – green triangle. Three independent experiments with similar results were pooled together for a total of 22 biological replicates for each gene. Error bars represent a 95% confidence interval.

To determine the membrane localization of SlySBP12a, the last 73 amino acids of the protein containing the putative TMD were fused to the C-terminal end of YFP (YFP-TMD_SlySBP12a_) (Supplementary Fig. S3). This construct was expressed in *N. benthamiana* where it localized to the periphery of epidermal cells and a ring-like structure around the nucleus that resembled endoplasmic reticulum (ER) localization (Fig. 5). Endoplasmic reticulum localization was confirmed in tomato protoplasts, where the YFP-TMD_SlySBP12a_ fusion colocalized with the ER marker SP-mCherry-HDEL (Fig. 7). This ER marker consists of the fluorescent protein mCherry with a signal peptide at its N-terminus and an ER retention motif at its C-terminus (Nelson *et al.*, 2007). We were also able to show colocalization between the full-length YFP-SlySBP12a construct and the ER marker in tomato protoplasts (Fig. 7). These results confirm that SlySBP12a contains a functional TMD that integrates the full-length protein into the ER membrane.

**Fig. 7.**
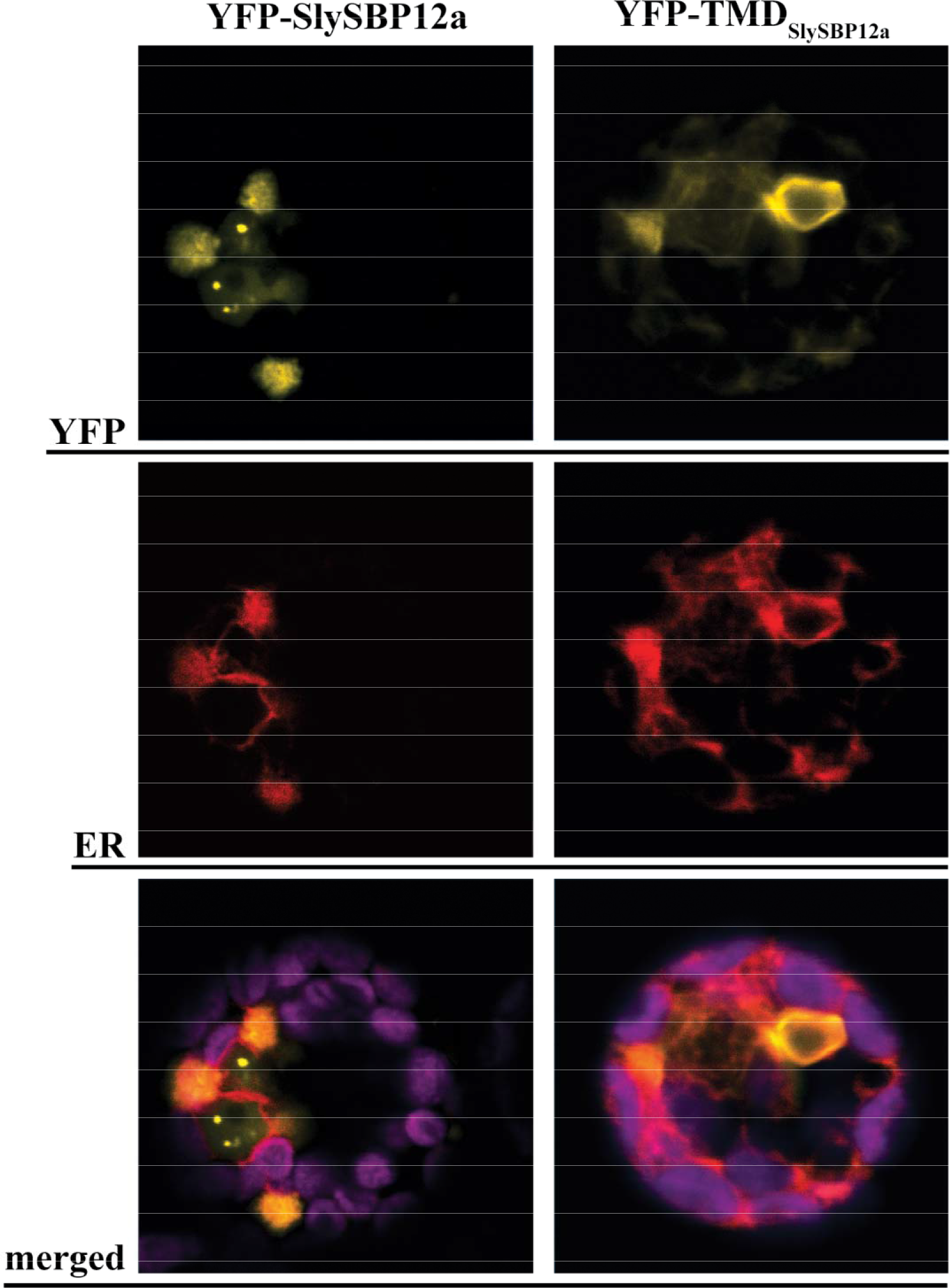
Endoplasmic reticulum localization of SlySBP12a and TMD_SlySBP12a_ in tomato protoplasts. Tomato protoplasts were transfected with plasmids encoding *35S:YFP-SiySBP12a* or *35S:YFP-TMD*_*SlySBP12a*_ and imaged by CLSM. An SP-mCherry-HDEL construct was co-transfected to serve as an ER marker (red). The magenta signal represents chloroplast autofluorescence.

### ROS production and fungal growth in leaves overexpressing SlySBP8b and SlySBP12a

Reactive oxygen species (ROS) are important cell death intermediaries, and their accumulation is a key feature of cell death imposed by necrotrophic fungal pathogens and the death-inducing toxins they produce (Heller and Tudzynski, 2011; Kim *et al.*, 2008; Sakamoto *et al.*, 2005; Shi *et al.*, 2007). Following transient expression in *N. benthamiana* leaves, we monitored the accumulation of hydrogen peroxide (H_2_O_2_) for four days using 3’3-diaminobenzidine (DAB) staining. Leaves expressing *35S:HA-SlySBP8b* and *35S:HA-SlySBP12a* displayed enhanced DAB staining intensity relative to expression of the *35S:YFP* control on the same leaf (Fig. 8A). Accumulation of H_2_O_2_ occurred as early as 2 and 3 days post-transformation for *35S:HA-SlySBP8b* and *35S:HA-SlySBP12a*, respectively (Fig. 8B). ImageJ software was used to measure DAB staining intensity (Schindelin *et al.*, 2012).

**Fig. 8.**
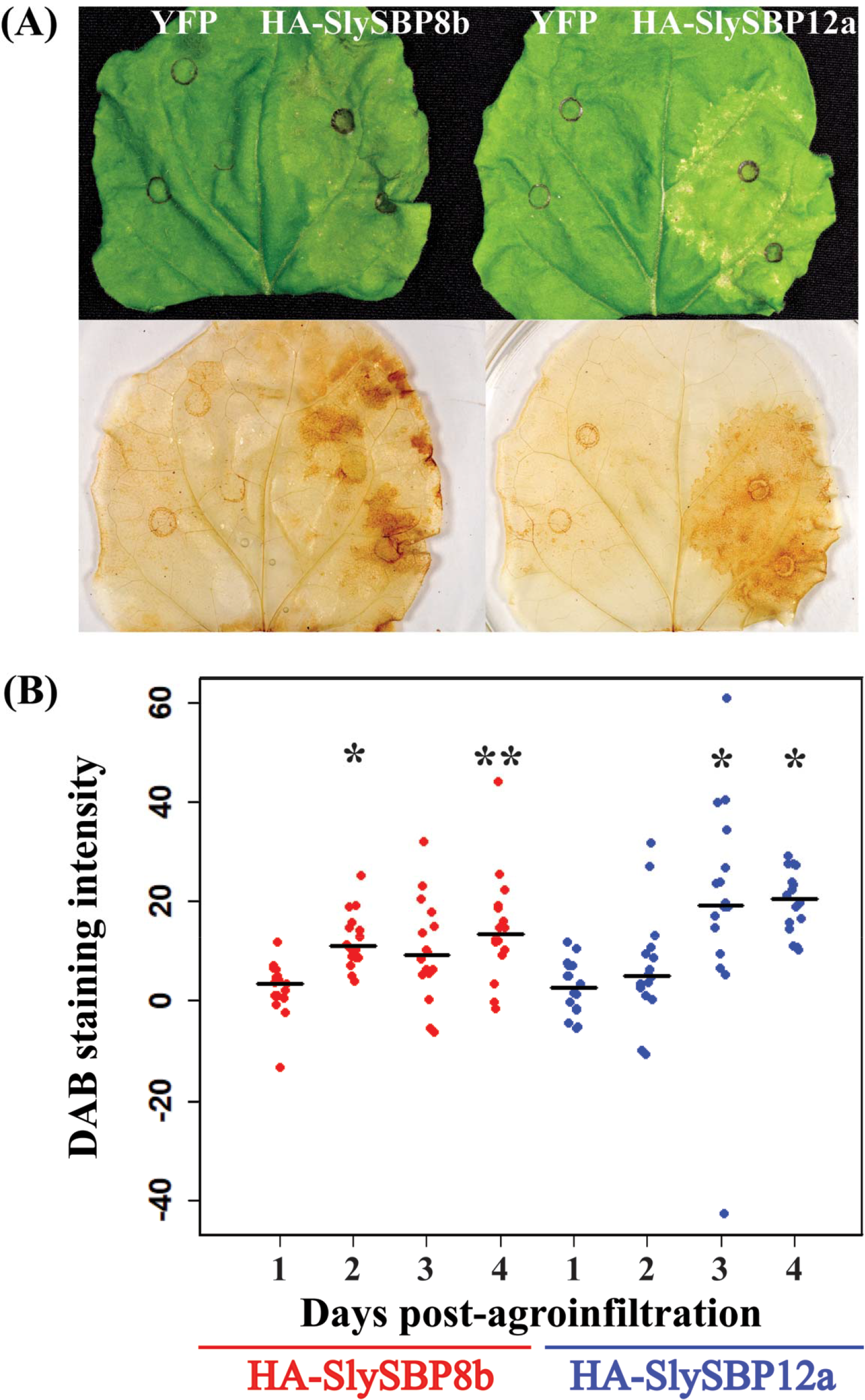
Overexpression of *SlySBP8b* and *SlySBP12a* in *N. benthamiana* induces H_2_O_2_ accumulation. *35S:HA-SlySBP8b* and *35S:HA-SlySBP12a* were transiently transformed in *N. benthamiana*. Leaves were cleared and stained with DAB to detect H_2_O_2_. (A) Images of leaves before and after DAB staining taken 4 days post-agroinfiltration. (B) Quantification of DAB-stained area for each *SlySBP* relative to *YFP* expression on the same leaf. ImageJ was used to analyze 16 leaves for each gene at each time point. All data points are displayed as a dotplot with the medians represented by black horizontal lines. Statistical significance compared to day 1 was determined using a one-way ANOVA with Tukey’s HSD post-hoc test (* P < 0.01; ** P < 0.001).

Transgenic *SfIAP* plants were reported to accumulate lower levels of ROS under stress conditions compared to wild-type plants (Li *et al.*, 2010). Necrotrophic fungal pathogens are known to exploit host ROS production as means to kill host cells for their own benefit (Govrin and Levine, 2000; Heller and Tudzynski, 2011; Kabbage *et al.*, 2013; Ranjan *et al.*, 2017). In addition to reduced ROS accumulation, transgenic *SfIAP* plants are also resistant to the necrotrophic fungal pathogen *A. alternata* (Li et al., 2010). Therefore, we reasoned that leaves overexpressing *SlySBP8b* and *SlySBP12a* would support enhanced growth of this pathogen. Unfortunately, *N. benthamiana* is not susceptible to this pathogen, so we screened *Nicotiana* germplasm for susceptible species (data not shown). We found that *Nicotiana glutinosa* was susceptible to *A. alternata* and previous work confirmed that transgenes could be expressed effectively in this species using Agrobacterium-mediated transient transformation (Kessens *et al.*, 2014).

While the differences were small, a total of 54 biological replicates from four randomized and blind experiments showed that leaves expressing *35S:YFP-SlySBP8b* or *35S:YFP-SlySBP12a* had increased *A. alternata* lesion areas compared to leaves expressing *35S:YFP* alone (Fig. 9A and 9B). This effect was more pronounced with *35S:YFP-SlySBP12a* than with *35S:YFP-SlySBP8b* expression. As a positive control, leaves were treated with 5 uM FB1 to simulate cell death induction by a fungal toxin. Lesion development in *35S:YFP-SlySBP12a*-expressing tissue and FB1-treated tissue was comparable (Fig. 9B). Fluorescence microscopy was used to confirm protein accumulation in each leaf before fungal inoculation and *35S:YFP-SlySBP8b* and *35S:YFP-SlySBP12a* were able to induce tissue death in *N. glutinosa* (data not shown). Lesion areas were measured using ImageJ software (Schindelin *et al.*, 2012). Overall, we show that these two transcription factors are able to increase ROS levels and promote *A. alternata* growth, phenotypes that are dampened in plants expressing *SfIAP*.

**Fig. 9.**
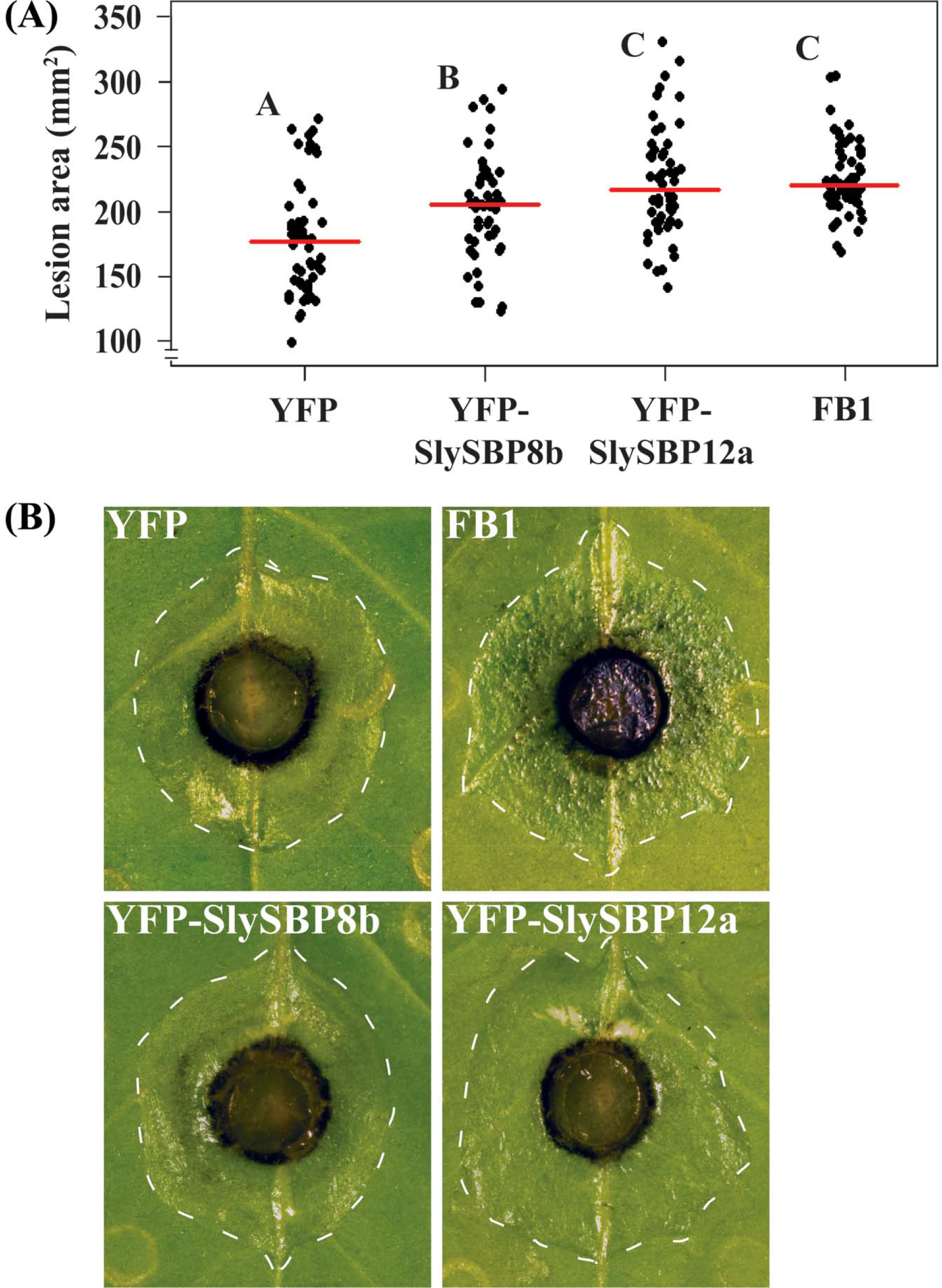
Overexpression of *SlySBP8b* and *SlySBP12a* enhances *A. alternata* growth on *N. glutinosa*. *35S:YFP-SlySBP8b*, *35S:YFP-SlySBP12a*, or *35S:YFP* were transiently transformed in *N. glutinosa*. As a positive control for cell death induction, leaves were treated with 5 μM FB1. Agar plugs containing actively growing *A. alternata* mycelium were placed fungal-side-down on leaves. (A) Quantification of lesion area using ImageJ. The results of 4 randomized and blind experiments were pooled representing 54 leaves for each treatment‥ All data points are displayed as a dotplot with the medians represented by red horizontal lines. Treatments with the same letter are not statistically significant as determined by a one-way ANOVA with Tukey’s HSD post-hoc test (YFP/SBP8b, P = 0.02; YFP/SBP12a, P = 2.0E−7; YFP/FB1, P = 5.0 E−7; SBP8b/SBP12a, P = 0.02; SBP8b/FB1, P = 0.04). (B) Images of inoculated leaves with lesions outlined by a dotted white line.

## Discussion

The first report of cell death suppression by heterologous expression of a viral IAP (*OpIAP*) in tobacco occurred almost two decades ago. *OpIAP* expression prevented cell death imposed by the necrotrophic fungal pathogen *S. sclerotiorum* and the necrosis-inducing viral pathogen tomato spotted wilt virus (Dickman *et al.*, 2001). Subsequent studies revealed that an IAP from *Spodoptera frugiperda* (SfIAP) suppressed cell death imposed by numerous abiotic and biotic stresses (Hoang *et al.*, 2014; Kabbage *et al.*, 2010; Li *et al.*, 2010). However, the biochemical mechanism by which these IAPs suppress cell death in plant systems remains unknown. In this study, we utilize SfIAP as a tool to identify novel pro-death regulators and provide a biochemical context for SfIAP function in plants.

### SlySBP8b and SlySBP12a associate with SfIAP^M4^(I332A) and exhibit characteristics of pro-death regulators

The yeast two-hybrid and CoIP data presented clearly show that SlySBP8b and SlySBP12a associate with SfIAP^M4^(I332A) (Table 1 & Fig. 2). Remarkably, *SlySBP8b* and *SlySBP12a* exhibit attributes of pro-death regulators, demonstrated by cell death induction and ROS accumulation upon overexpression (Fig. 1 and Fig. 8). We anticipated that coexpression of *SfIAP* with *SlySBP8b* or *SlySBP12a* would suppress cell death induction. However, numerous attempts to suppress cell death induced by *SlySBP8b* and *SlySBP12a* through *SfIAP* or *SfIAP*^*M4*^ coexpression were unsuccessful (data not shown). One possible explanation is the fact that SlySBP8b and SlySBP12a accumulate at higher levels compared to SfIAP and SfIAP^M4^, thus largely escaping SfIAP regulation. An excess of either transcription factor could allow enough to enter the nucleus and influence cell death gene expression.

SlySBP8b and SlySBP12a belong to a family of plant-specific transcription factors known as SQUAMOSA promoter binding proteins (SBPs), of which 15 members are present in tomato (Salinas *et al.*, 2012). Members of this family are defined by a highly conserved SBP-box DNA binding domain and can be further divided into 9 phylogenetically distinct clades (Preston and Hileman, 2013; Yamasaki *et al.*, 2013). *SBP* genes are known to regulate diverse developmental processes such as flowering time, branching, trichome development, apical dominance, and pollen sac development to name a few (Wang and Wang, 2015; Yamasaki *et al.*, 2013). Interestingly, silencing of the *SBP* gene *Colorless non-ripening* (*Cnr*) in tomato results in fruit with delayed ripening, a phenotype observed in tomatoes overexpressing *SfIAP* (Li *et al.*, 2010; Manning *et al.*, 2006).

While much is known about the role of SBP transcription factors in plant development, only a few studies to date have associated SBPs with stress responses. The deletion of Arabidopsis *SPL14* (*AtSPL14*) conferred enhanced tolerance to FB1, thus implicating this SBP transcription factor in the cellular response to this mycotoxin (Stone *et al.*, 2005). Tolerance to FB1 is a phenotype that we have also observed in *SfIAP*-overexpressing tomato seedlings (Li *et al.*, 2010). Interestingly, AtSPL14 and SlySBP12a both reside in clade-II and display similar structural characteristics with large SBP proteins that contain a predicted C-terminal transmembrane domain (Preston and Hileman, 2013).

Another clade-II member, *GmSPL12l* from soybean, was shown to be a target of the *Phakopsora pachyrhizi* (Asian soybean rust) effector PpEC23 (Qi *et al.*, 2016). This effector suppressed the hypersensitive response (HR) in soybean and tobacco and also interacted with other clade-II members from *N. benthamiana* and Arabidopsis: NbSPL1 and AtSPL1 (Qi *et al.*, 2016). In another study, the N immune receptor of *N. benthamiana* was found to associate with the SBP transcription factor NbSPL6 upon activation of HR. This interaction only occurred when plants were challenged with an HR-eliciting strain of tobacco mosaic virus (TMV-U1) but not a non-eliciting strain (TMV-Ob) (Padmanabhan *et al.*, 2013). Taken together, our results and the findings of previous studies clearly show that SBP transcription factors are critical regulators of plant stress responses that result in cell death.

Fungal pathogens with a necrotrophic lifestyle are known to exploit host ROS production for cell death induction and successful pathogenesis (Govrin and Levine, 2000; Heller and Tudzynski, 2011). As positive regulators of cell death and ROS production, we hypothesized that overexpression of *SlySBP8b* and *SlySBP12a* would support enhanced growth of necrotrophic fungal pathogens. Additionally, *SfIAP* transgenic plants are resistant to cell death induced by the necrotrophic fungal pathogen *A. alternata* (Li et al., 2010). The results of four randomized and blind experiments clearly show that while the contribution of *SlySBP8b* or *SlySBP12a* overexpression to *A. alternata* lesion areas was small, it was significantly greater than leaves expressing the negative control *35S:YFP* (Fig. 9). The small differences in growth could be explained by the fact that *A. alternata* is already an aggressive pathogen and the benefits of priming its host for death would be small. To test this, we also treated leaves with FB1, which is a structural analog of the AAL toxin produced by *A. alternata* f. sp. *lycopersici* that induces cell death in tomato (Mirocha *et al.*, 1992). Pre-treatment of *N. glutinosa* leaves with FB1 led to enhanced growth of *A. alternata* comparable to *SlySBP12a* overexpression (Fig. 9). These results provide further evidence that SlySBP8b and SlySBP12a are positive regulators of cell death, which in this case, contribute to pathogenic development of *A. alternata*.

### Nuclear localization of SlySBP8b and SlySBP12a is required for cell death

As members of a transcription factor family, we hypothesized that SlySBP8b and SlySBP12a exert their pro-death activity in the nucleus. We show that these transcription factors are clearly localized to the nucleus of tomato protoplasts (Fig. 3) and mutation of the bi-partite NLS of both transcription factors abolishes cell death (Fig. 4). These results support our hypothesis that SlySBP8b and SlySBP12a function in the nucleus, possibly through the regulation of genes involved in cell death. Future studies should focus on identifying the genes regulated by SlySBP8b and SlySBP12a, as this may provide further information on the downstream components responsible for cell death execution in plants.

### SlySBP12a localizes to the ER

Unlike SlySBP8b, which we found to be strictly nuclear localized, SlySBP12a was also present outside of the nucleus (Fig. 3 and Fig. 5). By fusing the putative C-terminal TMD of SlySBP12a to YFP, we were able to show that the TMD of SlySBP12a localized YFP around the nucleus and at the periphery of *N. benthamiana* epidermal cells (Fig. 5). We hypothesized that this pattern was due to ER localization. This was confirmed in tomato protoplasts, where both YFP-SlySBP12a and YFP-TMD_SlySBP12a_ co-localize with the ER marker SP-mCherry-HDEL (Fig. 7).

In response to environmental stress, plant cells increase production of secreted proteins, which in turn can cause ER stress due to the sudden influx of proteins that must be properly folded before moving through the rest of the secretory pathway (Eichmann and Schafer, 2012). This makes the ER an important sensor of cellular stress as the accumulation of unfolded proteins is first detected by the ER. Membrane-tethered transcription factors (MTTFs) residing at the ER membrane play important roles in ER stress perception and regulation of genes involved in stress relief and cell death in mammalian and plant systems (Slabaugh and Brandizzi, 2011). Membrane tethering provides spatial regulation of transcription factor activity, as MTTFs must be removed from the membrane before the transcription factor domain can translocate to the nucleus (Slabaugh and Brandizzi, 2011). This type of regulation allows these transcription factors to act quickly in response to cellular stress.

In this study, we show that SlySBP12a exhibits a localization pattern similar to previously described ER-MTTFs from Arabidopsis: NAC089, bZIP28, and bZIP60. These transcription factors are activated upon perception of ER stress and activate cell death (NAC089), heat stress (bZIP28), and ER stress (bZIP60) responses through transcriptional regulation of genes involved in these processes (Gao *et al.*, 2008; Iwata and Koizumi, 2005; Liu *et al.*, 2007; Yang *et al.*, 2014). Removal of the TMD from these transcription factors results in their complete localization to the nucleus and constitutive activation of the processes they regulate (Gao *et al.*, 2008; Iwata and Koizumi, 2005; Liu *et al.*, 2007; Yang *et al.*, 2014). This mirrors what we have observed with SlySBP12a. Removal of the TMD results in complete nuclear localization in *N. benthamiana* and tomato cells and enhanced cell death induction compared to full-length SlySBP12a (Fig. 3, Fig. 5, and Fig. 6).

With our data and previous studies of ER-MTTFs, we can speculate that SlySBP12a is cleaved from the ER membrane upon stress perception and translocates to the nucleus where it regulates genes involved in cell death. However, we must keep in mind that our experiments were performed with a cDNA copy of SlySBP12a, preventing the detection of splice-isoforms that could lack the TMD. This is important to consider as bZIP60 was originally thought to be proteolytically cleaved from the ER membrane upon stress induced by tunicamycin treatment (Iwata *et al.*, 2008). A follow-up study by the same group showed that in addition to being proteolytically cleaved, bZIP60 is also alternatively spliced in response to tunicamycin treatment, resulting in a truncated protein lacking the C-terminal TMD (Nagashima *et al.*, 2011). Future experiments looking at the translocation of SlySBP12a upon stress induction must consider the possibility of alternative splice isoforms.

## Conclusion

While the expression of *IAP* and other anti-apoptotic genes in plants confer enhanced stress tolerance, the animal-derived nature of these genes will likely prohibit their broad commercial use. Thus, the identification of endogenous plant cell death regulators, such as SBP transcription factors, that can be targeted to ameliorate stress tolerance is appealing. This is exemplified by recent interest in exploiting *SBP* genes for crop improvement due to the many developmental traits they regulate (Liu *et al.*, 2016; Wang and Wang, 2015). Efforts are underway in our lab to determine whether the disruption of these transcription factors impact tolerance to a range of abiotic and biotic insults.

## Supplementary Data

**Table S1.** List of primers used in this study.

**Fig. S1.** Summary of the QIS-Seq approach used to identify SfIAP-interacting partners.

**Fig. S2.** Western blot confirming SfIAP^M4^(I332A) is not cleaved at its N-terminus.

**Fig. S3.** Diagram of wild-type and mutant SlySBP8b and SlySBP12a constructs used in this study.

## Acknowledgements

We would like to thank Andrew Bent and members of his lab for their valuable insight and shared lab resources. The *Alternaria alternata* isolate used was provided by Shunping Ding of Amanda Gevens’ lab (University of Wisconsin-Madison). Next-generation sequencing for the yeast two-hybrid assay was performed by the Biotechnology Center at UW-Madison. Tomato protoplasts were generated with assistance from Stacy Anderson of Donna Fernandez’s lab (University of Wisconsin-Madison). Confocal microscopy was performed at the Newcomb Imaging Center at UW-Madison. *Nicotiana glutinosa* seeds were obtained from the U.S. Nicotiana Germplasm Collection donated by the University of California-Berkeley. R.K. is supported by a National Institutes of Health National Research Service Award T32 GM007215. This work was supported by a United States Department of Agriculture (USDA) Hatch award (WIS01818) to M.K.

